# Δ^9^-Tetrahydrocannabinol-induced enhancement of reward responsivity via mesocorticolimbic modulation in squirrel monkeys

**DOI:** 10.64898/2026.01.22.701118

**Authors:** Kwang-Hyun Hur, Lisa D. Nickerson, Jack Bergman, Jessi Stover, Stephen J. Kohut

## Abstract

Δ^9^-tetrahydrocannabinol (THC)-containing products are widely used recreationally, partly due to THC’s ability to enhance the appetitive (i.e., rewarding) properties of diverse stimuli. However, the neural mechanisms through which THC modulates reward-related processing remain poorly understood. Here, we used a Pavlovian paradigm in adult squirrel monkeys (3males, 1female) to associate a visual conditioned stimulus (CS^+^) with appetitive food delivery. The modulatory effects of acute THC (1-10μg/kg, i.m.) on behavioral and brain responses to CS^+^ were evaluated. Event-related functional MRI (fMRI) was employed to characterize the neural correlates of conditioned responding to the CS^+^, both in the absence and presence of THC treatment, with preconditioning scans serving as control. Behaviorally, THC (3μg/kg) selectively enhanced conditioned responding to the CS^+^ without altering responses to the control stimulus (CS^−^) or increasing baseline consummatory responding, underscoring the specificity of THC’s action on reward-associated processes. Consistently, fMRI analyses revealed that THC amplified CS^+^-evoked activation within mesocorticolimbic regions, including the anterior cingulate cortex (ACC), striatum, hippocampus, and substantia nigra-ventral tegmental area (SN-VTA), without affecting activity in visual and motor cortices. This finding underscores the selectivity of THC’s neuromodulatory effects on reward-related circuitry. Independent of CS exposure, resting-state functional connectivity analyses indicate that THC enhanced mesocorticolimbic network integration, as evident in strengthened SN-VTA-centered connectivity with the ACC, striatum, and hippocampus. Collectively, these findings demonstrate that THC enhances the responses to appetitive stimuli, through selective modulation of mesocorticolimbic circuitry, highlighting the SN-VTA as a pivotal hub for cannabinoid-mediated regulation of incentive salience and motivational drive toward reward-associated stimuli.

**One-sentence summaries:** THC enhances behavioral and neural responses to rewards through mesocorticolimbic modulation centered on the SN-VTA.

## 1. Introduction

Δ^9^-tetrahydrocannabinol (THC), the principal constituent of cannabis, is the most commonly used psychoactive substance worldwide [1]. Its increasing availability has raised significant public health concerns, including a higher prevalence of cannabis use disorder, associated adverse mental health outcomes, and rising hospitalization rates [2,3]. However, the abuse liability of THC remains controversial due to inconsistent clinical reports on its subjective psychoactive effects [4,5]. Furthermore, preclinical evidence for THC’s rewarding effects in conditioned place preference assays [6,7] or its reinforcing effects in self-administration studies [8-11] remains inconclusive. Of interest, the widespread co-use of THC with other substances raises the possibility that its primary action may not be as a rewarding or reinforcing stimulus but, instead, may be to enhance the rewarding properties of other stimuli [12]. Clinical findings provide some support for this hypothesis, indicating that longterm cannabis users often exhibit altered reward processing, characterized by heightened sensitivity to appetitive stimuli. In particular, chronic cannabis users demonstrate increased effort allocation for reward [13], as well as amplified neural activation in reward-related brain regions, including the striatum, in response to salient stimuli [14]. These effects of THC exposure are thought to contribute to its widespread use, underscoring the importance of further investigation to clarify THC’s potential reward-enhancing effects and its influence on motivational and neural responses to other salient stimuli.

THC’s reward-enhancing effects may be mediated by its pharmacological activity as a cannabinoid-1 (CB-1) receptor agonist [15], which modulates dopaminergic transmission within mesolimbic and mesocortical pathways, particularly those involving the substantia nigra-ventral tegmental area complex(SN-VTA) [16,17]. These dopaminergic pathways project to key forebrain structures such as the striatum, hippocampus, and prefrontal cortex, all of which are critically involved in motivational processes [18,19]. Despite these insights, the precise neurobiological mechanisms through which THC alters neural activity and functional connectivity within reward-related circuitry remain largely unexplored. In particular, investigations into the acute effects of THC on neural responses to reward-related stimuli are limited, as most existing neuroimaging studies have focused on chronic cannabis exposure [20,21]. Addressing this gap is essential for understanding how THC influences reward processing and for clarifying its role in the development and maintenance of substance use vulnerability.

Cue-reactivity studies employing functional magnetic resonance imaging (fMRI) have offered a valuable approach to characterize brain responses to reward-associated stimuli, and have enhanced our understanding of the neural mechanisms underlying reward-related disorders [22]. However, the ability to rigorously investigate these mechanisms in human subjects is constrained by ethical considerations and inherent sources of variability, including participant heterogeneity, uncontrolled substance use or psychiatric histories, and environmental confounds [23,24]. In this regard, translational fMRI employing standardized protocols in non-human primates (NHPs) allows rigorous control over experimental variables and thereby enhances the internal validity and predictive value of neuroimaging findings [25]. Critically, NHPs exhibit a high degree of neuroanatomical and functional homology with humans, especially within the mesocorticolimbic circuitry implicated in reward processing [19]. Such biological similarities provide a translationally relevant platform to investigate the neurobiological mechanisms that underlie reward-related responses.

The present study was conducted to elucidate the neurobiological mechanisms through which acute THC may modulate the behavioral and neural responses to appetitive stimuli (visual CS^+^) conditioned with palatable food delivery. In this work, a cue-reactivity fMRI paradigm in awake NHPs was employed to systematically determine how acute THC administration alters both regional activation and functional connectivity within reward-related circuitry, thereby providing mechanistic insight into its reward-enhancing effects.

## 2. Materials and Methods

### 2.1. Subjects

Four adult squirrel monkeys (*Saimiri sciureus*, three males and one female, Table.S1) were used, with each subject serving as its own control in a within-subject, repeated measures experimental design [26]. A power analysis based on relevant published studies from our laboratory and others demonstrated that a sample size of four monkeys provides sufficient statistical power to detect significant effects in both behavioral and neuroimaging studies [27-31]. Animal care and research procedures were conducted in accordance with approved protocols (see Supplement).

### 2.2. Drugs

THC was provided by the NIDA Drug Supply Program (Rockville, MD) and prepared in a vehicle comprising a 20:20:60 mixture of 95% ethanol, polysorbate-80 (Tween-80; Sigma-Aldrich, St Louis, MO), and 0.9% saline solution. Drug or vehicle injections were prepared in volumes of 0.3ml/kg body weight or less and administered intramuscularly (i.m.) 60-min prior to behavioral testing and MRI scanning.

#### 2.3.1. Pavlovian Conditioning

A Pavlovian conditioning paradigm [32] was employed to establish an association between a visual conditioned stimulus (CS) and highly palatable food reward(30% sweetened condensed milk; unconditioned stimulus, UCS) (Fig.1B). Subjects were seated in a prone posture within a custom-designed, MR-compatible chair that was placed inside a sound-attenuated and ventilated behavioral testing chamber [33]. A customized lickometer, which also served as the milk delivery apparatus, was positioned within tongue-reach of the subject. A light panel equipped with two distinct light-emitting diodes (LEDs; green on the left, red on the right) was placed 20cm in front of the subject’s face. One of the two lights served as the reward-paired stimulus (CS^+^), while the other served as the control stimulus (CS^−^), with the stimulus designation counterbalanced across subjects. Each session was comprised of three components and began with the illumination of a house light, signaling the onset of a 1,200-sec component composed of 40 discrete trials. In reward components, the CS^+^ was illuminated for 5-sec during each trial and 0.03ml of milk was delivered over 0.3-sec, simultaneous with the CS^+^ onset. In contrast, control components involved the illumination of the CS^−^ for 5-sec per trial without milk delivery. All trials in each component were followed by a 25-sec inter-trial interval (timeout). Reward components were conducted three times per day, while control components occurred once per day. The sequence of component types was counterbalanced across days and subjects to mitigate potential order effects. Training continued until the number of responses, measured as touches on the lickometer, to the CS^+^ in the absence of milk delivery exceeded that to the CS^−^ by at least five-fold, indicating successful acquisition of the conditioned association. All experimental procedures, including the presentation of stimuli, detection of responses, delivery of milk, and data acquisition, were controlled and recorded using Med Associates software (MedPC v4.2; Med Associates, St. Albans, VT).

**Figure 1.**
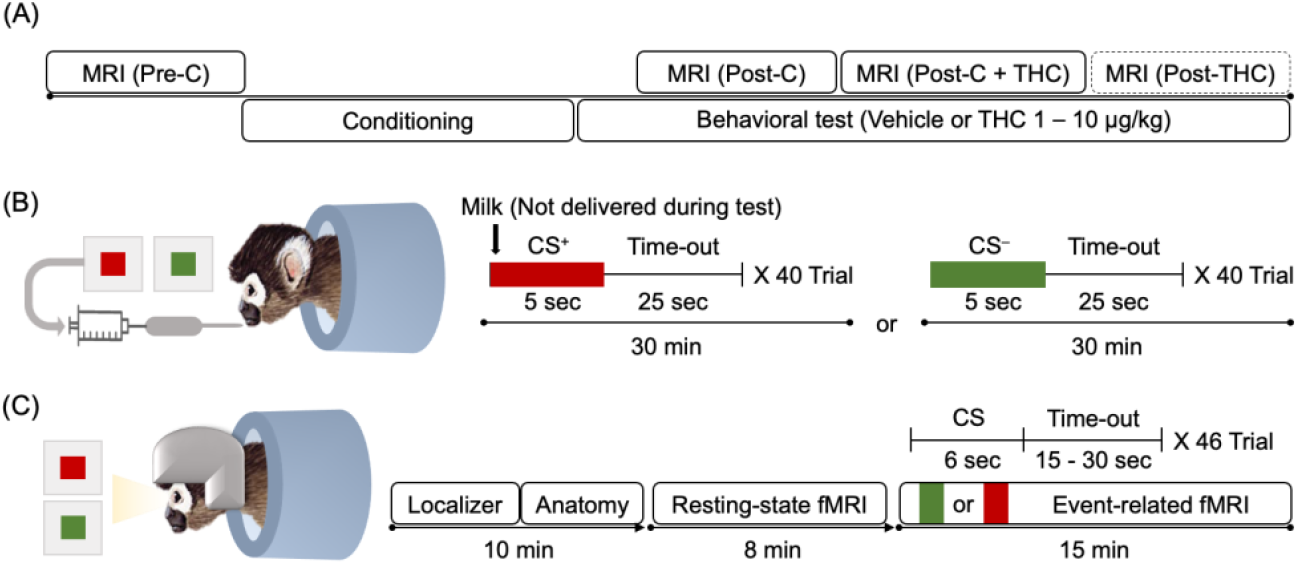
(A) Experimental timeline. Subjects (n = 4) underwent three main MRI sessions: Pre-conditioning (Pre-C), Post-conditioning (Post-C), and Post-conditioning with THC (Post-C+THC). An additional fMRI scan was conducted after the Post-C+THC session under the same conditions as the Post-C phase (referred to as Post-THC; see Supplementary). The Pre-C MRI was acquired prior to Pavlovian conditioning, during which a visual conditioned stimulus (CS) was paired with appetitive milk delivery. Following conditioning, behavioral responses to the CS were assessed under vehicle or THC administration (1–10 μg/kg) to determine the effective THC dose. Post-C and Post-C+THC MRI sessions were performed under vehicle or THC (3 μg/kg), respectively. Vehicle or THC was administered 60 min before behavioral testing and MRI acquisition. (B) Pavlovian conditioning. One of two colored lights (red or green) served as the reward-paired stimulus (CS^+^), while the other served as the control stimulus (CS^−^), with stimulus assignment counterbalanced across subjects. Each 1,200-sec CS session comprised 40 discrete trials. Each CS was illuminated for 5-sec, followed by a 25-sec time-out (inter-trial interval). Milk (0.03 mL/trial) was delivered concurrently with CS^+^, whereas CS^−^ trials were not reinforced. No milk was delivered during testing. (C) fMRI acquisition. Following anatomical scans, a resting-state fMRI scan was acquired, during which no CS were presented. Subsequently, event-related fMRI sessions were conducted, in which CS^+^ and CS^−^ were presented in a pseudorandom order. Each CS was presented for 6-sec, with jittered interstimulus intervals of 15-to 30-sec, resulting in a total of 23 presentations per CS. No milk was delivered during scanning.

#### 2.3.2. THC Effects on Conditioned Responding in the Absence of Milk Delivery

Following the completion of the conditioning phase, subjects underwent test sessions to evaluate the effects of various doses of THC (1-10μg/kg) on conditioned behavioral responses to the CS^+^ and CS^−^ in the absence of milk delivery. Behavioral measures included the total number of responses during the session and the temporal pattern of responding across the session. Each THC test session was separated by at least 7 days to minimize potential tolerance development. To maintain consistent conditioned responding across test sessions, behavioral training sessions continued on days between test sessions.

#### 2.3.3. THC Effects on Conditioned Responding During Milk Delivery and Food Consumption

To evaluate whether the effective dose of THC (3μg/kg) potentiated conditioned responding to the UCS (i.e. milk delivery), subjects were tested in experimental sessions in which CS^+^ presentations were paired with milk delivery. Two behavioral measures were analyzed: (1) the percentage of completed trials with a response occurring during the 5-second CS^+^ window, and (2) the response rate, defined as the number of responses per second during the CS^+^ session.

Subjects were tested further in their home cages to determine whether THC might enhance food consumption independent of conditioned stimuli. Following administration of vehicle or THC (3μg/kg), animals were provided with a food portion equivalent to 10% of their body weight. The time taken to consume the entire portion was recorded, and the consumption rate (g/min) was calculated for comparison across treatment conditions.

#### 2.4.1. Magnetic Resonance Imaging (MRI)

An event-related fMRI paradigm incorporating cue-reactivity [33], adapted from established clinical protocols [34,35], was employed to investigate brain responses to the CS in awake subjects (Fig.1C). During each fMRI session, reward-associated (CS^+^) and non-associated (CS^−^) visual stimuli were presented in a pseudorandom order to reduce anticipatory effects and maintain attentional engagement. Each CS was presented for 6-sec, with jittered interstimulus intervals ranging from 15-to 30-sec, totaling 23 presentations for each CS. These parameters were selected based on prior clinical cue-reactivity studies [36] and to match the expected hemodynamic response, which peaks at 5-to 8-sec and typically returns to baseline within 15-to 30-sec [37,38]. No rewards were delivered during scanning, enabling the isolated assessment of brain responses elicited by the CS.

Subjects underwent fMRI scans across three experimental phases (Fig.1A): pre-conditioning (Pre-C), post-conditioning (Post-C), and post-conditioning with THC (3μg/kg) administration (Post-C+THC). The THC dose (3μg/kg) used for fMRI was selected based on behavioral data showing that it reliably enhances conditioned responding to the reward-associated cue. Behavioral responses to the CS^+^ were confirmed prior to the Post-C and Post-C+THC fMRI sessions to ensure that conditioning remained intact. To control for potential confounding effects arising from continued conditioning over time, an additional fMRI scan was conducted following Post-C+THC session under the same conditions as the Post-C phase. Details on MRI data acquisition and preprocessing are provided in the Supplementary Information.

#### 2.3.4. Analysis of Brain Responses to CS

Whole-brain analysis was performed using FSL’s FEAT (FMRI Expert Analysis Tool) to examine brain (i.e. blood oxygen level-dependent, BOLD) responses to CS across three experimental phases: Pre-C, Post-C, and Post-C+THC. At the subject-level, event-related general linear models (GLMs) were constructed for each session, with separate regressors for CS^+^ and CS^−^ trials. Although each condition comprised 23 trials, to ensure the reliability of cue-reactivity estimates, only trials eliciting detectable BOLD responses in the primary visual cortex during CS presentation were retained. This inclusion criterion ensured that only visually attended trials contributed to the analysis. On average, 10.7±2 (SEM) trials per condition were included per subject. Stimulus events were convolved with a canonical hemodynamic response function (gamma function: phase=0, standard deviation=3, mean lag=6) to model expected BOLD responses. Temporal derivatives were included to account for variability in response timing, and motion-related noise was modeled using 12 degrees of freedom (6-motion parameters plus their temporal derivatives) as nuisance regressors. First-level contrasts were generated for CS^+^-activation, CS^+^-deactivation, CS^−^-activation, CS^−^-deactivation. These contrast images were carried forward for group-level analysis.

#### 2.4.3. Correlation between Behavioral and Brain Responses to CS^+^

To identify brain regions in which CS^+^-evoked brain activation was associated with behavioral responding, whole-brain correlation analyses were conducted using behavioral performance as a covariate within a repeated-measures group-level framework. For each subject, the behavioral measure was defined as the total number of responses during CS^+^ presentations in each experimental phase (Pre-C, Post-C, Post-C+THC). Behavioral data were demeaned across subjects prior to inclusion as a covariate in the GLM to ensure proper orthogonalization and facilitate the interpretability of covariate effects. Group-level analysis was conducted using mixed-effects analysis (FLAME 1+2) to account for within- and between-subject variance. Whole-brain statistical analysis was conducted using a voxel-wise threshold of p<0.001 (Z>3.1) without cluster-based correction for exploratory purposes, given the limited sample size, which may reduce sensitivity under stringent correction procedures. To balance sensitivity and specificity, only clusters exceeding 20 contiguous voxels were considered significant, thereby minimizing false positives while maintaining sufficient detection power [39]. Brain regions exhibiting significant positive or negative correlations between CS^+^-evoked BOLD activation and behavioral response were identified. These regions were delineated based on a standardized squirrel monkey brain atlas [40] and independently verified by two experienced neuroscientists.

To further characterize how phase-dependent changes in brain activity correlated with behavioral performance, β-values (parameter estimates) were extracted from both the full extent of significant clusters and anatomically defined regions of interest (ROIs). ROIs were defined as 3D spherical masks (0.5 search radius) centered on atlas-based coordinates (Fig.3E), including the anterior cingulate cortex (ACC), striatum, SN-VTA, hippocampus, visual cortex, and motor cortex. To avoid potential circular analysis [41], no further inferential statistics were conducted on these extracted data. Instead, these results are presented as illustrative supplementary findings to visualize brain-behavior correlations (Fig.3B) and phase-dependent changes in activity within each ROI (Fig.3C-3D).

#### 2.4.4. Analysis of Functional Connectivity between ROIs

To assess the impact of THC administration on intrinsic brain network dynamics, resting-state functional connectivity (rsFC) analysis was conducted using the 480-sec baseline period prior to CS exposure. This time window was selected to isolate THC-induced changes in rsFC from any potential influence of CS exposure. An ROI-based analytical approach was implemented, in which BOLD time-series data were extracted from each ROI during the baseline period. Partial correlation coefficients were computed for all pairwise combinations of ROIs using custom Python scripts, resulting in a 6×6 rsFC matrix for each subject. Partial correlation better estimates direct functional connectivity between ROIs by accounting for the influence of other regions, thereby reducing confounds from indirect pathways and enabling more precise assessment of inter-regional interactions within complex neural circuits [42]. To facilitate parametric statistical analysis at the group level and to ensure the normality of the data distribution, all individual correlation coefficients were transformed to z-values with Fisher’s r-to-z transformation [43].

### 2.5. Statistical Analysis

Statistical analyses were conducted using Prism 9.0 (GraphPad Software, San Diego, CA). Behavioral data were analyzed using repeated-measures ANOVA followed by Tukey’s post hoc test (Fig.2A and 2D) or using one-tailed paired t-tests for pairwise comparisons (Fig.2C and 2E). fMRI data were analyzed as described above using FSL’s FEAT (Fig.3A, Table.1A). Partial correlation coefficients derived from ROI-to-ROI rsFC analyses were compared between the Post-C and Post-C+THC phases using multiple paired t-tests (Fig.4B, Table.1B) and corrections for multiple comparisons were not applied in this case to avoid inflating Type II error rates, given the limited sample size and the associated reduction in statistical power. Statistical significance was defined as *p<0.05, **p<0.01.

**Figure 2.**
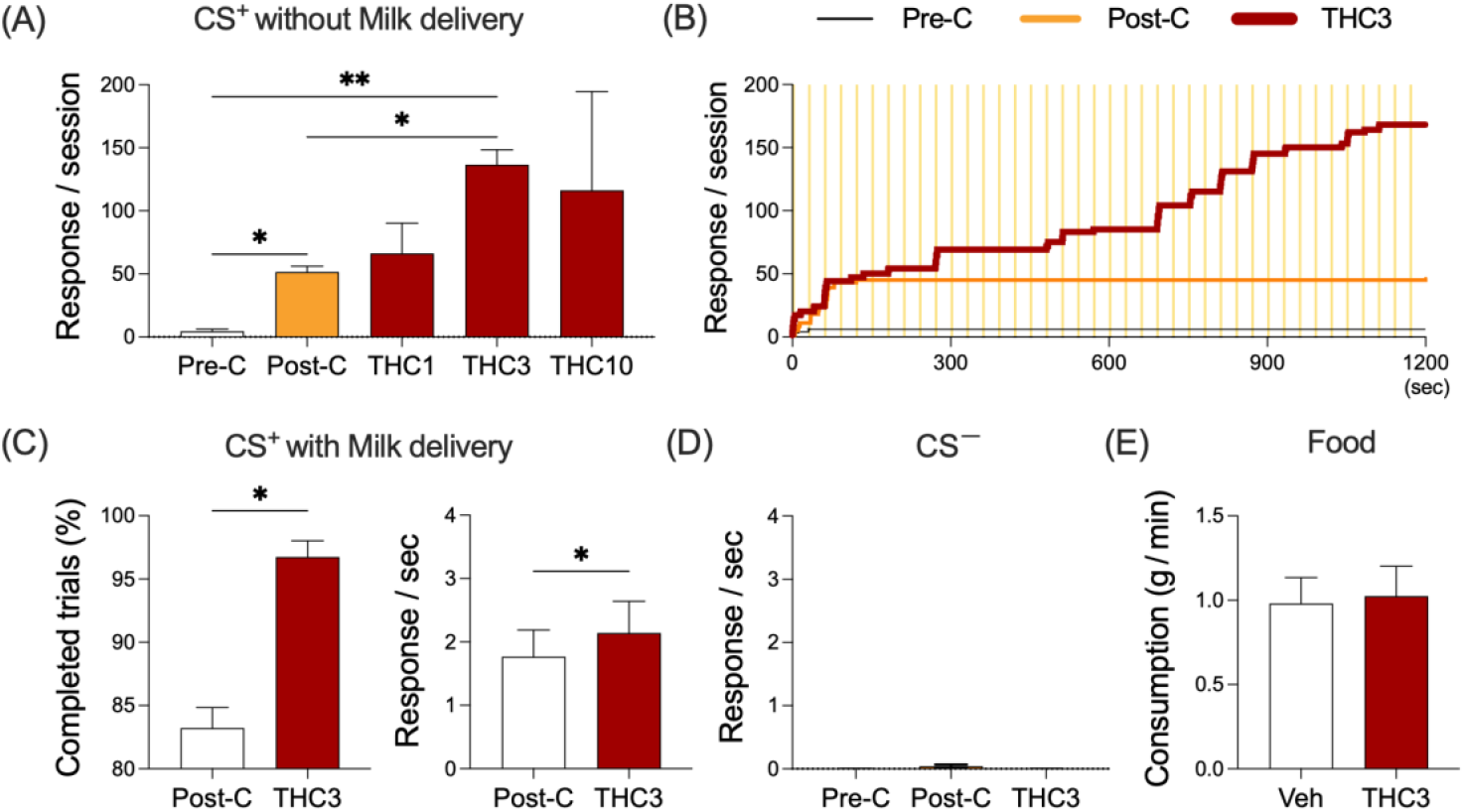
Effects of THC on conditioned behavioral responses to CS. Subjects (n = 4) underwent Pavlovian conditioning in which a reward-paired stimulus (CS^+^) was associated with milk delivery (0.03 mL/trial), while the control stimulus (CS^−^) remained unpaired. (A) Total number of responses during CS^+^ sessions without milk delivery across experimental phases: Pre-conditioning (Pre-C), Post-conditioning (Post-C), and Post-conditioning with THC (1-10 μg/kg, Post-C+THC). (B) Temporal distribution of responses during CS^+^ sessions across the Pre-C, Post-C, and Post-C+THC (3 μg/kg) phases from a representative subject. (C) Percentage of completed trials and response rate during CS^+^ sessions with milk delivery (D) Response rate during CS^−^ sessions across experimental phases. (F) Food consumption rate (g/min) following vehicle or THC (3 μg/kg) administration in the home cage. Data are presented as mean ± SEM. *p < 0.05, **p < 0.01.

**Figure 3.**
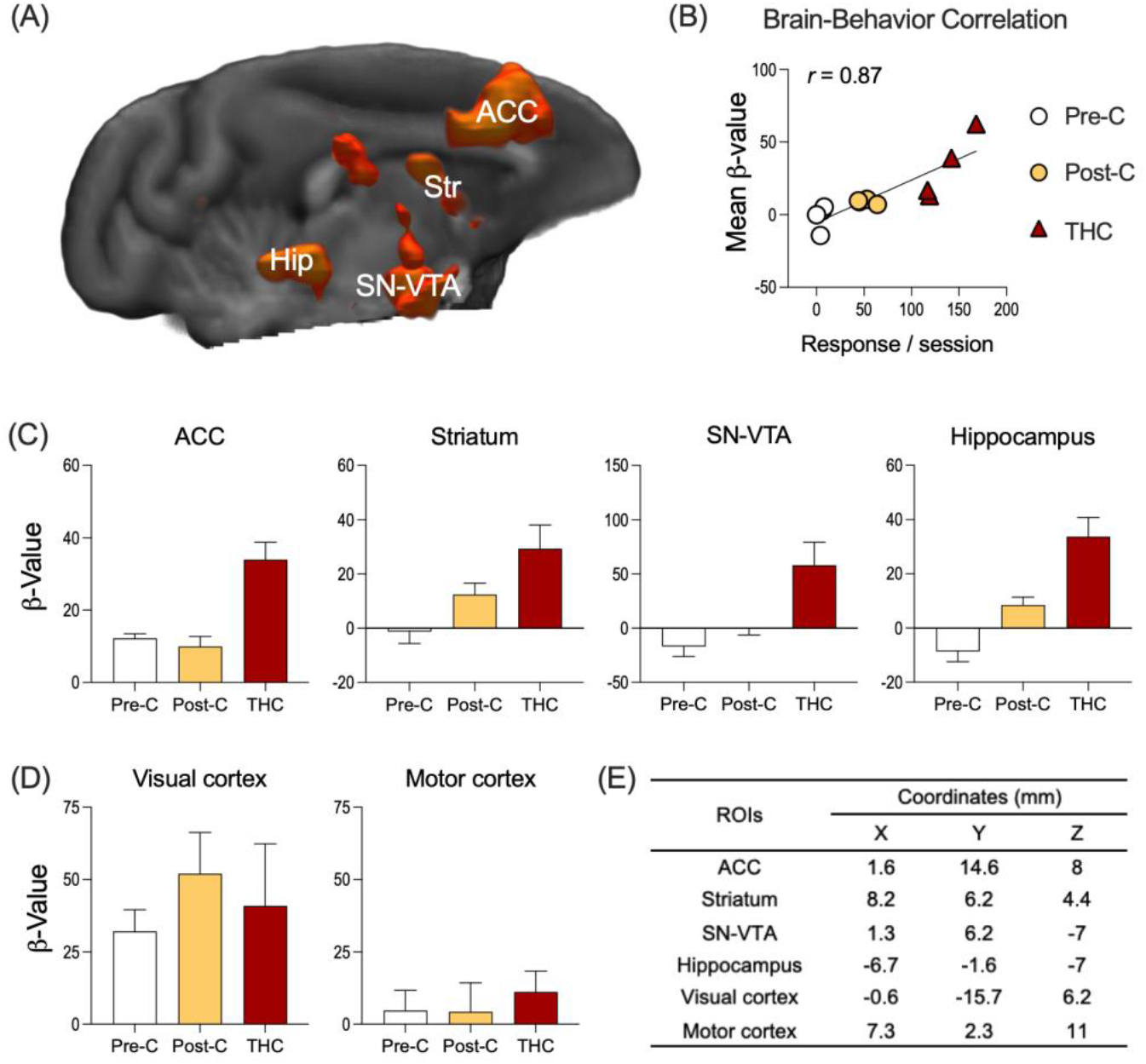
Brain regions exhibiting BOLD activation positively correlated with CS^+^-driven behavioral responses. Subjects (n = 4) underwent fMRI scanning across three experimental phases: Pre-C, Post-C, and Post-C+THC. (A) Whole-brain correlation analysis revealed brain regions showing significant positive associations between CS^+^-evoked BOLD activation and behavioral performance (i.e., total number of responses during CS^+^ sessions) across experimental phases. Significant clusters are spline-interpolated and overlaid onto a standardized 3D anatomical template of the squirrel monkey brain (right hemisphere view). Statistical maps are displayed using a red–yellow color scale (Z = 3.1-3.9). Identified regions include the anterior cingulate cortex (ACC), striatum (Str), hippocampus (Hip), and substantia nigra–ventral tegmental area (SN-VTA). (B) Correlation between mean β-values extracted from the full extent of significant clusters and behavioral performance across experimental phases is shown for visualization purposes. Each data point represents an individual subject at each phase. (C-D) Phase-dependent changes in CS^+^-evoked BOLD activation were examined in ROIs identified by the correlation analysis, as well as in control ROIs, including the visual and motor cortices. β-values were extracted from each ROI to illustrate the effects across the three experimental phases. No statistical analyses were conducted on these data to avoid potential circular analysis (e.g., as seen in (B), the group effect is embedded in the behavioral association). (E) Atlas-based coordinates of each ROI.

**Table 1.**

| Correlation | Atlas Coordinates (mm) |  |  |  |  |  | Brain regions |
| --- | --- | --- | --- | --- | --- | --- | --- |
|  | Cluster | # Voxels | Z-Max | X | Y | Z |  |
| Positive | 11 | 403 | 5.58 | 2.29 | 15.2 | 10.9 | Anterior cingulate cortex |
|  | 10 | 317 | 5.68 | -7.65 | -0.694 | 4.31 | Parietal area_L, Temporal area_L |
|  | 9 | 211 | 6.05 | 2.29 | 5.27 | -6.92 | Substantia nigra, Ventral tegmental area |
|  | 8 | 120 | 5.56 | 5.27 | -6.66 | -4.11 | Hippocampus_R |
|  | 7 | 100 | 5.48 | -6.66 | -2.68 | -6.92 | Hippocampus_L |
|  | 6 | 86 | 4.94 | -7.65 | -8.64 | -10.7 | Cerebellum_L |
|  | 5 | 70 | 4.41 | 6.26 | -0.694 | 6.18 | Parietal area_R |
|  | 4 | 49 | 4.58 | 8.25 | 6.26 | 4.31 | Striatum_R |
|  | 3 | 47 | 4.43 | 11.2 | 5.27 | -4.11 | Temporal_R |
|  | 2 | 40 | 4.57 | -7.65 | 10.2 | -1.31 | Striatum_L |
|  | 1 | 38 | 4.11 | 1.29 | 12.2 | -0.371 | Striatum_R |
| Negative | N/A |  |  |  |  |  |  |

| Brain Regions | | Partial correlation (z-score, Mean ± SEM) | | Differences | t-ratio | p-value |
| --- | --- | --- | --- | --- | --- | --- |
|  |  | Post-C | Post-C+THC |  |  |  |
| SN-VTA | ACC | 0.19 ± 0.12 | 0.36 ± 0.12 | 0.17 ± 0.01 | 12.37 | **< 0.01\*\*** |
| | Str | 0.05 ± 0.06 | 0.22 ± 0.1 | 0.17 ± 0.04 | 4.68 | **0.02\*** |
| | Hip | 0.21 ± 0.03 | 0.48 ± 0.06 | 0.27 ± 0.08 | 3.30 | **0.04\*** |
| | VC | $0.02 \pm 0.08$ | $0.01 \pm 0.1$ | $0 \pm 0.08$ | 0.06 | 0.96 |
| | MC | $0.05 \pm 0.09$ | $0.05 \pm 0.05$ | $0 \pm 0.11$ | 0.02 | 0.99 |
| ACC | Str | $0.16 \pm 0.09$ | $0.08 \pm 0.09$ | $-0.07 \pm 0.13$ | 0.59 | 0.59 |
| | Hip | $0.12 \pm 0.07$ | $0.12 \pm 0.03$ | $0 \pm 0.05$ | 0.03 | 0.98 |
| | VC | $0.15 \pm 0.08$ | $0.18 \pm 0.09$ | $0.03 \pm 0.14$ | 0.21 | 0.84 |
| | MC | $0.11 \pm 0.09$ | $0.03 \pm 0.06$ | $-0.08 \pm 0.08$ | 1.00 | 0.39 |
| Str | Hip | $0.11 \pm 0.07$ | $0.14 \pm 0.07$ | $0.03 \pm 0.13$ | 0.24 | 0.83 |
| | VC | $0.21 \pm 0.15$ | $0.19 \pm 0.1$ | $-0.03 \pm 0.08$ | 0.33 | 0.76 |
| | MC | $-0.04 \pm 0.11$ | $-0.04 \pm 0.07$ | $0 \pm 0.04$ | 0.09 | 0.93 |
| Hip | VC | $0.07 \pm 0.05$ | $0.1 \pm 0.08$ | $0.03 \pm 0.12$ | 0.22 | 0.84 |
| | MC | $0.1 \pm 0.04$ | $0.06 \pm 0.14$ | $-0.03 \pm 0.17$ | 0.19 | 0.86 |
| VC | MC | $0.07 \pm 0.14$ | $0.09 \pm 0.14$ | $0.02 \pm 0.12$ | 0.16 | 0.88 |

## 3. Results

### 3.1. Selective enhancement of reward-associated behaviors following THC treatment

When the CS^+^ was presented in the absence of milk delivery (Fig.2A), subjects exhibited minimal responding during the Pre-C phase (Mean±SEM: 4.5±1.7). In contrast, a significant increase in conditioned responding was observed during the Post-C phase (51.5±4.62; p=0.01, q=11.43), indicating successful acquisition of the Pavlovian association. Administration of 3 μg/kg THC further enhanced conditioned responding (136.5±11.93), which was significantly greater than in both the Pre-C (p<0.01, q=16.16) and Post-C (p = 0.04, q = 7.61) phases. Specifically, THC administration led to sustained responding to CS^+^ presentations (Fig.2B and S1), with a significant increase in the number of responses during the latter half of test sessions (Table.S2).

THC (3μg/kg) treatment significantly increased the percentage of completed trials during sessions in which the CS^+^ was presented with food delivery (Fig.2C; 96.75±1.27%) compared to the Post-C phase 83.23±1.62% (p=0.01, t=5.38). Likewise, the response rate during CS^+^ sessions was significantly elevated under the THC condition (2.14±0.49) relative to the Post-C phase (1.76±0.42; p=0.02, t=4.5).

In contrast to its enhancement of responding to the CS^+^, no significant differences in response rate were noted during CS^−^ sessions across experimental phases (Fig.2D; Pre-C, Post-C, Post-C+THC; p=0.33, f=0.15), indicating the specificity of the THC effect to reward-paired CS. Additionally, the rate of food consumption in the home cage did not significantly differ between vehicle and THC (3μg/kg) conditions (Fig.2E; p=0.22, t=0.92), suggesting that the doses of THC in the present study did not alter consummatory behavior.

### 3.2. Brain regions showing correlation between CS^+^-evoked BOLD activation and behavioral responses

Fig.3A illustrates brain regions (details in Table.1A), including the ACC, striatum, SN-VTA, and hippocampus, in which significant positive correlations between CS^+^-evoked BOLD activity and behavioral measures (i.e., total number of responses during CS^+^ sessions) were observed, with Fig.3B providing a visual representation of these correlations across experimental phases (Pre-C, Post-C, Post-C+THC).

Fig.3C and 3D visually depicts changes in CS^+^-evoked BOLD activity in ROIs across experimental conditions. In the ACC, activity did not significantly differ between the Pre-C (12.2±1.29) and Post-C (9.99±2.72) phases but increased significantly in the Post-C+THC condition (34±4.76). In the striatum, SN-VTA, and hippocampus, activity significantly increased following conditioning and was further enhanced by THC administration: Striatum, Pre-C (−1.22±4.43), Post-C (12.42±4.21), Post-C+THC (29.31±8.72); SN-VTA, Pre-C (−16.96±9.15), Post-C (−0.29±5.99), Post-C+THC (58.01±21.22); Hippocampus, Pre-C (−8.64±3.79), Post-C (8.47±2.89), Post-C+THC (33.68±7.12). Activity in the visual cortex increased in response to conditioning (Pre-C, 32.08±7.52; Post-C, 52.02±14.25) but was not further modulated by THC (40.93±21.44). In contrast, the motor cortex did not exhibit significant changes in activity across conditioning or THC administration: Pre-C (4.72±6.98), Post-C (4.25±10.03), Post-C+THC (11.13±7.23).

### 3.3. Effects of THC on rsFC between ROIs

Differences in ROI-to-ROI rsFC between the Post-C and Post-C+THC phases are illustrated in Fig. 4A. Notably, THC administration induced significant increases in rsFC between the SN-VTA and several ROIs, including the ACC (p<0.01, t=12.37), striatum (p=0.01, t=4.68), and hippocampus (p=0.04, t=3.29) (Fig.4B). In contrast, no significant changes in rsFC were observed between other ROI pairs across the two experimental phases (Table.1B).

**Figure 4.**
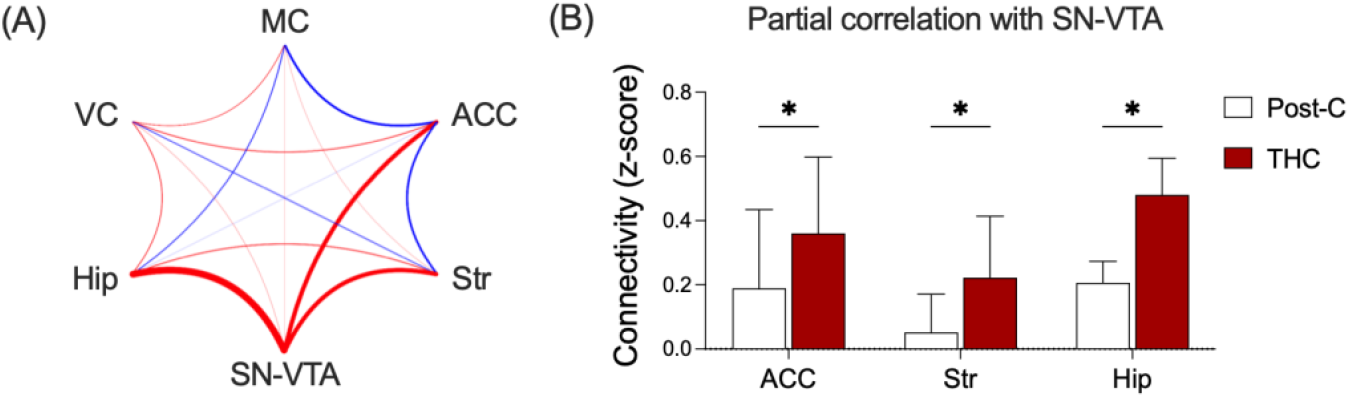
THC-induced alterations in rsFC between brain regions. ROI-to-ROI rsFC was evaluated during the baseline period preceding CS exposure in the Post-C and Post-C+THC phases (n = 4). ROIs included the anterior cingulate cortex (ACC), striatum (Str), hippocampus (Hip), substantia nigra–ventral tegmental area (SN-VTA), visual cortex (VC), and motor cortex (MC). (A) Connectome diagram depicting differences in ROI-to-ROI rsFC between the Post-C and Post-C+THC phases. Line thickness reflects the magnitude of connectivity change; red lines indicate increased connectivity (Post-C+THC > Post-C), and blue lines indicate decreased connectivity (Post-C+THC < Post-C). (B) Phase-dependent differences in rsFC between the SN-VTA and each of the following ROIs: ACC, Str, and Hip. Data are presented as mean ± SEM. *p < 0.05.

## 4. Discussion

The present study employed a cue-reactivity fMRI paradigm in awake NHPs to investigate the neurobiological mechanisms by which acute THC administration may modulate responses to appetitively conditioned stimuli. Behavioral results show that the acute administration of a relatively low dose of THC (3μg/kg) produced enhancement of responding to a visual stimulus (CS^+^), that was previously conditioned to the delivery of highly palatable milk (UCS), under both milk-absent and milk-present conditions. These findings suggest that THC enhances the attribution of incentive salience to reward-predictive cues under non-reinforced conditions, while also enhancing motivational engagement during reinforced conditions. Such effects align with clinical evidence indicating that cannabis use is associated with heightened responsiveness to reward-associated stimuli [14] and increased behavioral effort to secure reinforcement [13]. Notably, the THC-induced enhancement observed in the present study was absent at a higher dose (10μg/kg), highlighting the dose-selective nature of THC’s effects on motivated behavior and suggesting that the reward-enhancing effects at a low dose (3μg/kg) were unlikely to be confounded by non-specific behavioral impairments typically induced by higher doses of the cannabinoid [44]. Importantly, THC had no impact on responses to CS^−^ or on unconditioned food consumption in the home cage, further indicating that its behavioral effects selectively target conditioned motivational processes rather than general motor or consummatory functions.

Whole-brain correlation analyses between CS^+^-evoked BOLD responses and behavioral performance revealed significant engagement of mesocorticolimbic circuitry, including the ACC, striatum, SN-VTA, and hippocampus, in response to appetitive conditioning and THC administration. The trajectory of BOLD activity (i.e., β-values) across experimental phases in each ROI illustrated how brain activation was modulated by both conditioning and THC treatment. The striatum and SN-VTA, core components of the midbrain dopaminergic system, exhibited robust activation following conditioning, reflecting their critical involvement in reinforcement learning [45]. This activation was further enhanced by THC administration, in line with previous evidence that cannabinoids enhance midbrain activity [46], thereby promoting the encoding of reward-associated stimuli [47]. The hippocampus also showed significant conditioning-related activation, consistent with its well-recognized role in encoding and retrieval of contextual memory [48]. THC administration augmented hippocampal engagement, potentially facilitating the sustained retrieval of the CS^+^-reward association and thereby maintaining behavioral responsiveness to the CS^+^. Conversely, the ACC did not exhibit robust activation following conditioning alone but showed significant engagement following THC treatment. Considering the ACC’s established role in evaluating reward salience and regulating goal-directed behavior [49], such findings indicate that THC could increase the motivational value of reward-associated stimuli, thereby enhancing behavioral responses to the CS^+^ in the present study. Notably, THC administration did not elicit significant changes in BOLD responses within visual or motor cortices, underscoring the regional specificity of its effects. Taken together, these findings indicate that low doses of THC may preferentially modulate neural circuits involved in reward processing while sparing primary sensory and motor functions [44].

To provide complementary insights into phase-dependent changes in responses to CS^+^ and CS^−^, which served as a control condition, a whole-brain ANOVA was performed across experimental phases (see Supplementary Methods.1.3 and Table.S3). Regarding brain responses to the CS^+^, relative to the Pre-C phase, the Post-C phase revealed enhanced activation in the visual cortex, as well as parietal and temporal areas, which are implicated in visuospatial attention [50,51], likely reflecting the use of visual stimuli during conditioning. However, these visual-related activations were not further augmented following THC administration (Post-C vs. Post-C+THC); instead, THC selectively induced significant activation in reward-related brain regions, including the ACC and striatum. Comparisons between Pre-C and Post-C+THC phases revealed concurrent increases in both visual and reward-related regions, providing converging evidence with the correlation analyses. In contrast, for CS^−^, conditioning elicited activation in the visual cortex (Pre-C<Post-C), whereas no THC-induced enhancement was observed in reward-related brain regions. Overall, these findings demonstrate that THC selectively potentiates the response to conditioned reward-predictive cues by modulating activity within reward-related circuits.

Furthermore, given that the Post-C+THC session was conducted subsequent to the Post-C session, an additional fMRI scan (referred to as Post-THC) was performed under the same conditions as the Post-C phase to evaluate potential confounding effects arising from the additional conditioning sessions (See Supplementary Methods.1.4). Comparative analyses revealed no significant increases in CS^+^-evoked brain activation within mesocorticolimbic circuits during the Post-THC session relative to the Post-C phase (Table.S4), accompanied by no significant differences in behavioral responding across sessions (Table.S5). These findings indicate that the observed THC-induced enhancement in brain and behavioral responses to the CS^+^ is specifically attributable to the acute pharmacological effects of THC, rather than to extended conditioning or task repetition.

rsFC analyses of mesocorticolimbic circuitry revealed that THC selectively increased connectivity between the SN-VTA, the primary dopaminergic hub, and key mesocorticolimbic regions, including the ACC, striatum, and hippocampus. Such effects support the notion that THC primarily enhances reward-related neural signaling within these circuits. These findings are consistent with the ability of cannabinoids to both facilitate dopamine release in the SN-VTA and enhance firing of dopaminergic neurons projecting to limbic and cortical targets [46,52], and further align with previous studies showing that mesocorticolimbic rsFC strength depends on dopaminergic neurotransmission, with stronger dopamine signaling associated with greater coupling between SN-VTA and its projection regions [53]. Importantly, no significant changes in rsFC were observed among other ROI pairs in the present study, suggesting that acute THC may selectively potentiate the dopaminergic signaling from the SN-VTA to its cortical and limbic projection targets. Overall, consistent with evidence that increased mesocorticolimbic rsFC correlates with heightened brain responses to reward-associated stimuli [54], these findings suggest that THC selectively strengthens functional synchrony within reward-related neural networks, thereby enhancing incentive salience of reward-predictive cues (CS^+^).

Several limitations to the present study should be acknowledged. First, although the sample size was sufficient for within-subject comparisons and neuroimaging analyses, the relatively small cohort constrained the statistical power to detect potential subtle effects. Second, the present investigation found that a low dose of THC (3μg/kg) was sufficient to selectively enhance motivational salience without eliciting widespread behavioral disruption. However, future studies utilizing a broader dose range, including higher doses (e.g., 10μg/kg), are warranted to delineate the dose-dependent transition from facilitative to disruptive effects of THC on reward processing and associated neural circuits. Lastly, our cue-reactivity paradigm used a food-reward associative framework and the generalizability of the present effects to other reward modalities (e.g., drug-related, social rewards) remains uncertain. Future studies incorporating larger sample sizes, systematic dose-response manipulations, and a broader range of stimulus categories will be essential for determining the robustness, dose sensitivity, and cross-modal applicability of THC’s effects on reward processing. Additionally, longitudinal designs could help clarify whether acute THC-induced modulation of mesocorticolimbic circuits predicts persistent changes in motivational bias or susceptibility to reward-related disorders.

In conclusion, this study provides novel evidence that acute THC administration selectively modulates functional activity and connectivity within mesocorticolimbic dopaminergic circuits, thereby enhancing behavioral and neural responses to appetitively conditioned stimuli. These findings elucidate mechanistic pathways through which THC potentiates motivational processes and suggest that its popularity may, in part, reflect an amplification of the rewarding properties of other stimuli. Importantly, understanding THC’s impact on reward processing is critical for clarifying its contribution to the development and maintenance of substance use vulnerability, and it may accelerate the translation of these insights into targeted preventive strategies for THC-related interventions. Furthermore, our awake NHP cue-reactivity neuroimaging model provides a highly translational platform for studying reward processing, allowing real-time measurement of neural circuit dynamics in response to reward-associated stimuli. This methodological advance may facilitate investigations into the neurobiological mechanisms underlying reward-related disorders characterized by abnormal reward sensitivity.

## Acknowledgments

This research was supported by grants from the Joseph & Susan Gatto Foundation Award (KHH) and NIH/NIDA R01 DA048150 (JB and SJK). The authors thank Julia Cunningham and Marissa Costa for expert technical assistance in conducting experiments.

## Author Contribution

Kwang-Hyun Hur: Conceptualization, Investigation, Methodology, Formal analysis, Writing - Original Draft, Visualization, Funding acquisition Lisa D. Nickerson: Formal analysis Jack Bergman: Conceptualization, Supervision, Funding acquisition, Writing - Review & Editing, Jessi Stover: Investigation Stephen J. Kohut: Conceptualization, Methodology, Resources, Writing - Review & Editing, Supervision, Funding acquisition

## Data and Materials Availability

All data needed to evaluate the conclusions in the paper are present in the paper. Raw data from this study are available from the corresponding author upon reasonable request.

## References

1. UNODC, World Drug Report 2023. United Nations publication, 2023.

2. Di Forti, M., et al., The contribution of cannabis use to variation in the incidence of psychotic disorder across Europe (EU-GEI): a multicentre case-control study. The Lancet Psychiatry, 2019. 6(5): p. 427–436.

3. Hall, W., et al., Public health implications of legalising the production and sale of cannabis for medicinal and recreational use. The Lancet, 2019. 394(10208): p. 1580–1590.

4. Metrik, J., et al., Acute effects of marijuana smoking on negative and positive affect. Journal of Cognitive Psychotherapy, 2011. 25(1): p. 31–46.

5. Crane, N.A. and K.L. Phan, Effect of Δ9-Tetrahydrocannabinol on frontostriatal resting state functional connectivity and subjective euphoric response in healthy young adults. Drug and alcohol dependence, 2021. 221: p. 108565.

6. Moklas, M.A.M., et al., Chronic delta-9-tetrahydrocannabinol induces monoamine release but not conditioned place preference. The Open Behavioral Science Journal, 2012. 6(1).

7. Moore, C.F., et al., Δ9-Tetrahydrocannabinol Vapor Exposure Produces Conditioned Place Preference in Male and Female Rats. Cannabis and Cannabinoid Research, 2022.

8. Justinova, Z., et al., Self-administration of Δ 9-tetrahydrocannabinol (THC) by drug naive squirrel monkeys. Psychopharmacology, 2003. 169: p. 135–140.

9. Stringfield, S.J. and M.M. Torregrossa, Intravenous self-administration of delta-9-THC in adolescent rats produces long-lasting alterations in behavior and receptor protein expression. Psychopharmacology, 2021. 238: p. 305–319.

10. Leite, J.R. and E. Carlini, Failure obtain “cannabis-directed behavior” and abstinence syndrome in rats chronically treated with Cannabis sativa extracts. Psychopharmacologia, 1974. 36(2): p. 133–145.

11. Mansbach, R., et al., Failure of Delta (9)-tetrahydrocannabinol and CP 55,940 to maintain intravenous self-administration under a fixed-interval schedule in rhesus monkeys. Behavioural pharmacology, 1994. 5(2): p. 219–225.

12. Maksimovic, A. and H. Heacock, Co-use of cannabis with commonly used licit and illicit drugs. BCIT Environmental Public Health Journal, 2019.

13. Vele, K.C., J.M. Cavalli, and A. Cservenka, Effort-based decision making and self-reported apathy in frequent cannabis users and healthy controls: a replication and extension. Journal of Clinical and Experimental Neuropsychology, 2022. 44(2): p. 146–162.

14. Jager, G., et al., Tentative evidence for striatal hyperactivity in adolescent cannabis-using boys: a crosssectional multicenter fMRI study. Journal of psychoactive drugs, 2013. 45(2): p. 156–167.

15. Pertwee, R.G., Ligands that target cannabinoid receptors in the brain: from THC to anandamide and beyond. Addiction biology, 2008. 13(2): p. 147–159.

16. French, E.D., Δ9-Tetrahydrocannabinol excites rat VTA dopamine neurons through activation of cannabinoid CB1 but not opioid receptors. Neuroscience letters, 1997. 226(3): p. 159–162.

17. Bossong, M.G., et al., Δ9-tetrahydrocannabinol induces dopamine release in the human striatum. Neuropsychopharmacology, 2009. 34(3): p. 759–766.

18. Wise, R.A., Dopamine, learning and motivation. Nature reviews neuroscience, 2004. 5(6): p. 483–494.

19. Haber, S.N. and B. Knutson, The reward circuit: linking primate anatomy and human imaging. Neuropsychopharmacology, 2010. 35(1): p. 4–26.

20. Kangas, B.D., et al., Chronic Δ9-tetrahydrocannabinol exposure in adolescent nonhuman primates: persistent abnormalities in economic demand and brain functional connectivity. Neuropsychopharmacology, 2025. 50(3): p. 576–585.

21. Kohut, S.J., et al., Effects of cannabinoid exposure on short-term memory and medial orbitofrontal cortex function and chemistry in adolescent female rhesus macaques. Frontiers in Neuroscience, 2022. 16: p. 998351.

22. Childress, A.R., et al., Cue reactivity and cue reactivity interventions in drug dependence. NIDA research monograph, 1993. 137: p. 73–73.

23. Saccà, L., The uncontrolled clinical trial: scientific, ethical, and practical reasons for being. Internal and emergency medicine, 2010. 5: p. 201–204.

24. Hurst, S.A., et al., Ethical difficulties in clinical practice: experiences of European doctors. Journal of medical ethics, 2007. 33(1): p. 51–57.

25. Howell, L.L. and K.S. Murnane, Nonhuman primate neuroimaging and the neurobiology of psychostimulant addiction. Annals of the New York Academy of Sciences, 2008. 1141(1): p. 176–194.

26. Lamb, G.D., Understanding” within” versus” between” ANOVA Designs: Benefits and Requirements of Repeated Measures. 2003.

27. Adewale, A.S., D.M. Platt, and R.D. Spealman, Pharmacological stimulation of group ii metabotropic glutamate receptors reduces cocaine self-administration and cocaine-induced reinstatement of drug seeking in squirrel monkeys. Journal of Pharmacology and Experimental Therapeutics, 2006. 318(2): p. 922–931.

28. Mishra, A., et al., Functional connectivity with cortical depth assessed by resting state fMRI of subregions of S1 in squirrel monkeys. Human brain mapping, 2019. 40(1): p. 329–339.

29. Goense, J.B. and N.K. Logothetis, Neurophysiology of the BOLD fMRI signal in awake monkeys. Current Biology, 2008. 18(9): p. 631–640.

30. Grech, D.M., R.D. Spealman, and J. Bergman, Self-administration of D 1 receptor agonists by squirrel monkeys. Psychopharmacology, 1996. 125: p. 97–104.

31. Withey, S.L., et al., Fentanyl-induced changes in brain activity in awake nonhuman primates at 9.4 Tesla. Brain Imaging and Behavior, 2022. 16(4): p. 1684–1694.

32. Berridge, K.C., Reward learning: Reinforcement, incentives, and expectations, in Psychology of learning and motivation. 2000, Elsevier. p. 223–278.

33. Yassin, W., et al., Resting-state networks of awake adolescent and adult squirrel monkeys using ultrahigh field (9.4 T) functional magnetic resonance imaging. Eneuro, 2024. 11(5).

34. Gold, A.L., et al., Age differences in the neural correlates of anxiety disorders: An fMRI study of response to learned threat. American Journal of Psychiatry, 2020. 177(5): p. 454–463.

35. Britton, J.C., et al., Response to learned threat: An fMRI study in adolescent and adult anxiety. American Journal of Psychiatry, 2013. 170(10): p. 1195–1204.

36. Schlauch, R.C., et al., Psychometric evaluation of the Substance Use Risk Profile Scale (SURPS) in an inpatient sample of substance users using cue-reactivity methodology. Journal of psychopathology and behavioral assessment, 2015. 37: p. 231–246.

37. Katwal, S.B., et al., Measuring relative timings of brain activities using fMRI. NeuroImage, 2013. 66: p. 436–448.

38. Kim, J.H., et al., Dynamics of the cerebral blood flow response to brief neural activity in human visual cortex. Journal of Cerebral Blood Flow & Metabolism, 2020. 40(9): p. 1823–1837.

39. Fusar-Poli, P., Voxel-wise meta-analysis of fMRI studies in patients at clinical high risk for psychosis. Journal of Psychiatry and Neuroscience, 2012. 37(2): p. 106–112.

40. Gergen, J.A. and P.D. MacLean, A stereotaxic atlas of the squirrel monkey’s brain (Saimiri sciureus). 1962: US Department of Health, Education, and Welfare, Public Health Service ….

41. Kriegeskorte, N., et al., Circular analysis in systems neuroscience: the dangers of double dipping. Nature neuroscience, 2009. 12(5): p. 535–540.

42. Marrelec, G., et al., Partial correlation for functional brain interactivity investigation in functional MRI. Neuroimage, 2006. 32(1): p. 228–237.

43. Van Dijk, K.R., et al., Intrinsic functional connectivity as a tool for human connectomics: theory, properties, and optimization. Journal of neurophysiology, 2010. 103(1): p. 297–321.

44. Katsidoni, V., A. Kastellakis, and G. Panagis, Biphasic effects of Δ9-tetrahydrocannabinol on brain stimulation reward and motor activity. International journal of neuropsychopharmacology, 2013. 16(10): p. 2273–2284.

45. Cox, J. and I.B. Witten, Striatal circuits for reward learning and decision-making. Nature Reviews Neuroscience, 2019. 20(8): p. 482–494.

46. French, E.D., K. Dillon, and X. Wu, Cannabinoids excite dopamine neurons in the ventral tegmentum and substantia nigra. Neuroreport, 1997. 8(3): p. 649–652.

47. Luján, M.Á., et al., Mobilization of endocannabinoids by midbrain dopamine neurons is required for the encoding of reward prediction. Nature communications, 2023. 14(1): p. 7545.

48. Xia, L., et al., Dorsal-CA1 hippocampal neuronal ensembles encode nicotine-reward contextual associations. Cell reports, 2017. 19(10): p. 2143–2156.

49. Shenhav, A., M.M. Botvinick, and J.D. Cohen, The expected value of control: an integrative theory of anterior cingulate cortex function. Neuron, 2013. 79(2): p. 217–240.

50. Silver, M.A., D. Ress, and D.J. Heeger, Topographic maps of visual spatial attention in human parietal cortex. Journal of neurophysiology, 2005. 94(2): p. 1358–1371.

51. Ramezanpour, H. and M. Fallah, The role of temporal cortex in the control of attention. Current Research in Neurobiology, 2022. 3: p. 100038.

52. Bloomfield, M.A., et al., The effects of Δ9-tetrahydrocannabinol on the dopamine system. Nature, 2016. 539(7629): p. 369–377.

53. Bär, K.-J., et al., Functional connectivity and network analysis of midbrain and brainstem nuclei. Neuroimage, 2016. 134: p. 53–63.

54. Adrián-Ventura, J., et al., Reward network connectivity “at rest” is associated with reward sensitivity in healthy adults: A resting-state fMRI study. Cognitive, Affective, & Behavioral Neuroscience, 2019. 19(3): p. 726–736.

