## Supplementary Materials for "Δ^9^-Tetrahydrocannabinol-induced enhancement of reward responsivity via mesocorticolimbic modulation in squirrel monkeys"

### 1. Supplementary Methods

#### 1.1. Subjects

Subjects were housed in stainless-steel cages within a temperature- and humidity-controlled vivarium, with a 12-hour light/dark cycle (lights on 07:00-19:00). Monkeys were maintained at approximate free-feeding weights through daily feedings of a nutritionally balanced diet comprised of high-protein chow (#5040, LabDiet, St. Louis, MO) along with fresh fruits and vegetables. Water was available *ad libitum* outside of the experimental session. An assortment of manipulable toys, mirrors, puzzles, and background music during light hours were provided as part of an environmental enrichment program. Animal care and research were conducted according to the guidelines provided by the Institute of Laboratory Animal Resources and the National Institutes of Health Office of Laboratory Animal Welfare. The facility is licensed by the U.S. Department of Agriculture, and all experimental protocols were approved by the Institutional Animal Care and Use Committee at McLean Hospital.

#### 1.2. MRI Data Acquisition and Preprocessing

All MRI data were acquired using a 9.4 Tesla horizontal bore magnet system (400-mm diameter; Varian Inc., Palo Alto, CA). The system was equipped with an 11.6-cm inner-diameter gradient coil capable of producing a maximum gradient strength of 45 G/cm. A single-loop transmit-receive RF surface coil, optimized to encompass the subjects' head, was used for signal reception transmit and receive. Image localization and automated shimming were performed prior to data acquisition to ensure field homogeneity. Whole-brain fMRI data were acquired using a gradient-echo EPI sequence with the following parameters: echo time (TE)=8-ms, repetition time (TR)=1,500-ms, flip angle=90°, voxel resolution=0.99×0.99×0.93-mm, scan matrix=64×64, and field of view (FOV)=64-mm. The protocol included 54 coronal slices with a thickness of 1-mm, and 1,200 volumes were collected over a 30-min acquisition period. For anatomical reference and spatial normalization, distortion-matched structural images were acquired using a spin-echo EPI sequence with the following parameters: TE=17.5-ms, TR=1,500-ms, flip angle=90°, 8 signal averages, scan matrix=64×64, and FOV=64-mm. The anatomical dataset consisted of 54 coronal slices (1-mm thickness), precisely aligned to match the functional imaging slices. Throughout MRI scanning sessions, subjects were monitored in real-time via a live video feed (12M camera, MRC Systems GmbH, Heidelberg, Germany). Physiological monitoring of heart rate and blood oxygen saturation ($SpO_2$) was performed using the Nonin 7500 pulse oximeter (Nonin Medical Inc., Plymouth, MN), with measurements recorded at 5-min intervals to ensure subjects' safety.

Data were visually inspected for artifacts, and quality control was performed using the MRI Quality Control tool[1]. Intensity spikes were identified and corrected (https://github.com/bbfrederick/spikefix) with a threshold of 1-mm framewise displacement. Preprocessing was carried out using FMRIB's Software Library (FSL, Oxford University, UK), including the removal of the first 10 volumes from each scan to allow for data stabilization. Motion correction was performed using the MCFLIRT tool in FSL[2]. Spatial smoothing was applied using a Gaussian kernel with a 2.0-mm full-width at half-maximum (FWHM), and temporal filtering was performed with a high-pass filter at a 100-sec cutoff (0.01-Hz). Session-averaged functional volumes were aligned to the VALiDATe T2-weighted template[3] using a 12 degrees of freedom (DOF) affine transformation, followed by adjustment for nonlinear distortion fields using the JIP analysis toolkit (www.nitrc.org/projects/jip). Skull-stripping was applied to isolate brain tissue from non-brain structures.

#### 1.3. Comparison of Brain Responses to CS across Experimental Phases

To evaluate phase-dependent changes in brain activity, whole-brain ANOVA was performed using FSL's FEAT (FMRI Expert Analysis Tool) to identify differences in blood oxygen level-dependent (BOLD) responses to conditioned stimuli (CS) across three experimental phases: pre-conditioning (Pre-C), post-conditioning (Post-C), and post-conditioning with THC administration (Post-C+THC). Subject-level statistical maps were generated using event-related general linear models (GLMs) with separate regressors for $CS^+$ and $CS^-$ trials. Contrast images representing $CS^+$- and $CS^-$-evoked neural activity were then entered into a higher-level mixed-effects model (FLAME 1+2), which accounted for both within- and between-subject variability. Group-level analysis employed GLM-based ANOVA to assess phase-specific differences in neural responses to CS. Whole-brain statistical analysis was conducted using a voxel-wise threshold of $p < 0.001$ ($Z > 3.1$) without cluster-based correction for exploratory purposes, given the limited sample size, which may reduce sensitivity under stringent correction procedures. To balance sensitivity and specificity, only clusters exceeding 20 contiguous voxels were considered significant, thereby minimizing false positives while maintaining sufficient detection power [4]. Brain regions involved in the identified significant clusters were delineated using a standardized squirrel monkey brain atlas [5] and independently verified by two experienced neuroscientists. Detailed results are presented in Table S3.

#### 1.4. Control for Extended Conditioning Effects

To control for potential confounding effects arising from continued Pavlovian conditioning over time, an additional fMRI scan was conducted following Post-C+THC session under the same conditions as the Post-C phase (referred to as Post-THC). This design allowed for direct comparison between the Post-C and Post-THC phases to dissociate THC-specific neural effects from those attributable to extended training or task repetition. Event-related GLMs were again constructed at the subject level with separate regressors for $CS^+$ and $CS^-$ trials. First-level contrast maps were carried forward to group-level mixed-effects analysis (FLAME 1+2). Paired t-tests were used to assess phase-specific changes in neural responses between Post-C and Post-THC phases. Statistical inference followed the same thresholding and correction procedures as described above. Significant clusters were anatomically defined using the squirrel monkey brain atlas [5] and verified by two experienced neuroscientists. Detailed results are presented in Table S4.

**References**


1. Esteban, O., et al., *MRIQC: Advancing the automatic prediction of image quality in MRI from unseen sites.* PloS one, 2017. **12**(9): p. e0184661.
2. Jenkinson, M., et al., *Improved optimization for the robust and accurate linear registration and motion correction of brain images.* Neuroimage, 2002. **17**(2): p. 825-841.
3. Schilling, K.G., et al., *The VALiDATe29 MRI based multi-channel atlas of the squirrel monkey brain.* Neuroinformatics, 2017. **15**: p. 321-331.
4. Fusar-Poli, P., *Voxel-wise meta-analysis of fMRI studies in patients at clinical high risk for psychosis.* Journal of Psychiatry and Neuroscience, 2012. **37**(2): p. 106-112.
5. Gergen, J.A. and P.D. MacLean, *A stereotaxic atlas of the squirrel monkey's brain (Saimiri sciureus)*. 1962: US Department of Health, Education, and Welfare, Public Health Service ….

### 3. Supplementary Figures

**Figure S1.** Temporal distribution of behavioral responses during CS$^+$ sessions across all subjects in the Pre-C, Post-C, and Post-C+THC (3 µg/kg) phases. Each session lasted 1200-sec and included 40 CS$^+$ trials without reward delivery.

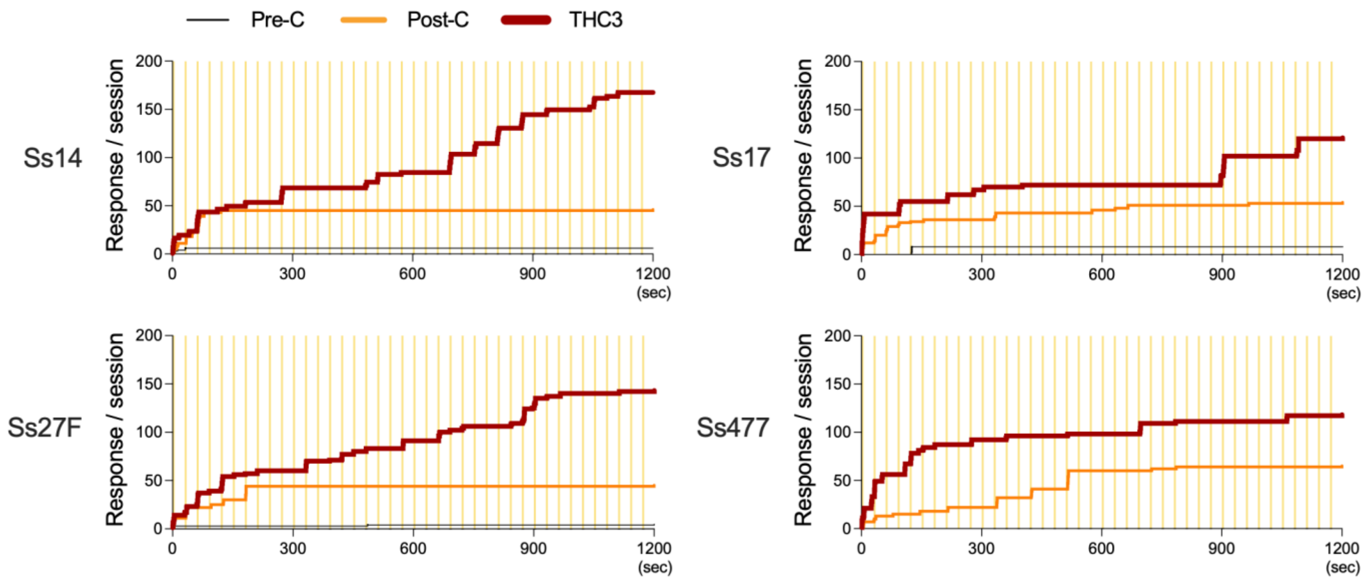

### 4. Supplementary Tables

**Table S1.** Characteristics of each monkey

| Subjects ID | Sex | Age (Years) | Weight (g) |
|---|---|---|---|
| Ss14 | Male | 5 | 800 |
| Ss17 | Male | 5 | 650 |
| Ss27F | Female | 5 | 650 |
| Ss477 | Male | 15 | 1000 |

**Table S2.** Number of responses during early and late $CS^+$ sessions, compared between Post-Conditioning (Post-C) and Post-Conditioning with THC administration (Post-C+THC) phases. Pairwise comparisons were conducted using one-tailed paired t-tests. Statistical significance was set at *p < 0.05, **p < 0.01.

| **$CS^+$ sessions** | Number of responses (Mean ± SEM) | | Differences | t-ratio | p-value |
|---|---|---|---|---|---|
| | Post-C | Post-C+THC | | | |
| Early (0 – 600 sec) | 48.75 ± 3.77 | 86.5 ± 5.52 | 37.75 ± 4.36 | 8.65 | < 0.01** |
| Late (600 – 1200 sec) | 2.75 ± 1.7 | 50.25 ± 13.09 | 47.5 ± 14.06 | 3.38 | 0.02* |

**Table S3.** Brain regions showing significant differences in CS-evoked BOLD responses across the three experimental phases (Pre-C, Post-C, and Post-C+THC). Whole-brain analyses were conducted using GLMs within a repeated-measures ANOVA framework to assess phase-dependent changes in neural responses to each CS ($CS^+$ and $CS^-$). Clusters comprising more than 20 voxels ($p < 0.001$, uncorrected) are listed, with anatomical localization determined using a standardized squirrel monkey brain atlas. R = right; L = left.

| **$CS^+$ Activation** | Cluster | # Voxels | Z-Max | Atlas Coordinates (mm) | | | Brain regions |
|---|---|---|---|---|---|---|---|
| | | | | X | Y | Z | |
| Pre-C < Post-C | 6 | 115 | 4.42 | -15.6 | -3.68 | -2.24 | Temporal area_L |
| | 5 | 110 | 3.83 | 2.29 | -19.6 | 6.18 | Visual cortex |
| | 4 | 73 | 4.25 | 10.2 | 4.27 | -4.11 | Temporal area_R |
| | 3 | 37 | 3.94 | 9.24 | -13.6 | 10.9 | Parietal area_R |
| | 2 | 30 | 3.8 | 8.25 | -2.68 | -5.05 | Cerebellum_R |

| | Cluster | # Voxels | Z-Max | X | Y | Z | Brain regions |
|---|---|---|---|---|---|---|---|
| | 1 | 26 | 4.46 | -2.68 | -6.66 | -5.05 | Cerebellum_L |
| Pre-C > Post-C | | | | N/A | | | |
| Post-C < Post-C+THC | 4 | 57 | 4.06 | -1.69 | 15.2 | 10.9 | Anterior cingulate cortex |
| | 3 | 46 | 4.93 | 4.27 | 9.24 | 7.11 | Striatum_R |
| | 2 | 24 | 3.53 | 6.26 | 6.26 | -10.7 | Amygdala_R |
| | 1 | 21 | 3.71 | 6.26 | -14.6 | -11.6 | Cerebellum_R |
| Post-C > Post-C+THC | | | | N/A | | | |
| Pre-C < Post-C+THC | 14 | 702 | 5.47 | -8.64 | -2.68 | 4.31 | Parietal area_L, Temporal area_L, Hippocampus_L, Cerebellum_L |
| | 13 | 307 | 5.3 | 12.2 | 4.27 | -5.98 | Temporal area_R, Substantia nigra, Ventral tegmental area |
| | 12 | 270 | 5.49 | 4.27 | 9.24 | 6.18 | Anterior cingulate cortex, Striatum_R |
| | 11 | 179 | 5.49 | 5.27 | -5.66 | 7.11 | Parietal area_R |
| | 10 | 157 | 4.56 | -7.65 | -25.5 | -1.31 | Visual cortex |
| | 9 | 146 | 6.05 | 7.26 | -4.67 | -5.98 | Cerebellum_R, Hippocampus_R |
| | 8 | 124 | 4.6 | 0.3 | -20.6 | 4.31 | Visual cortex |
| | 7 | 89 | 5.18 | 5.27 | -14.6 | -7.85 | Cerebellum_R |
| | 6 | 60 | 4.6 | 1.29 | -24.5 | -5.05 | Visual cortex |
| | 5 | 47 | 4.24 | 0.3 | -14.6 | 1.5 | Visual cortex |
| | 4 | 26 | 3.69 | 2.29 | 5.27 | 14.6 | Parietal area_R |
| | 3 | 24 | 3.7 | 5.27 | -25.5 | -0.371 | Visual cortex |
| | 2 | 23 | 4.43 | -0.694 | -18.6 | -9.72 | Visual cortex |
| | 1 | 22 | 3.87 | 8.25 | 5.27 | 5.24 | Parietal area_R |
| Pre-C > Post-C+THC | | | | N/A | | | |

| **CS⁻ Activation** | | | | Atlas Coordinates (mm) | | | Brain regions |
|---|---|---|---|---|---|---|---|
| | Cluster | # Voxels | Z-Max | X | Y | Z | |
| Pre-C < Post-C | 2 | 56 | 4.03 | -6.66 | -18.6 | -3.18 | Visual cortex_L |
| | 1 | 33 | 3.63 | 6.26 | -18.6 | 2.44 | Visual cortex_R |
| Pre-C > Post-C | | | | N/A | | | |
| Post-C < Post-C+THC | 1 | 50 | 4.23 | 1.29 | 19.2 | -0.371 | Orbitofrontal Cortex |
| Post-C > Post-C+THC | 1 | 25 | 4.57 | -8.64 | 7.26 | -4.11 | Amygdala_L |
| Post-C > Post-C+THC | | | | N/A | | | |
| Pre-C < Post-C+THC | 1 | 48 | 4.01 | 12.2 | -12.6 | -3.18 | Visual cortex_R |
| Pre-C > Post-C+THC | | | | N/A | | | |

**Table S4.** Brain regions exhibiting significant differences in CS-evoked BOLD responses between the Post-C and Post-THC phases. Whole-brain analyses were performed using GLMs within a paired t-test framework to identify phase-dependent changes in BOLD responses to each CS ($CS^+$ and $CS^-$). Anatomical localization of significant clusters ($p < 0.001$, uncorrected, >20 voxels) was determined using a standardized squirrel monkey brain atlas. R = right; L = left.

| **$CS^+$ Activation** | Atlas Coordinates (mm) | | | | | | Brain regions |
|---|---|---|---|---|---|---|---|
| | Cluster | # Voxels | Z-Max | X | Y | Z | |
| Post-C < Post-THC | 1 | 77 | 4.03 | -1.69 | -24.5 | -6.92 | Visual cortex |
| Post-C > Post-THC | 6 | 81 | 4.25 | 4.27 | 16.2 | 12.7 | Dorsolateral prefrontal cortex_R |
| | 5 | 60 | 4.23 | 8.25 | 13.2 | 6.18 | Ventrolateral prefrontal cortex_R |
| | 4 | 58 | 4.06 | -7.65 | 13.2 | 3.37 | Striatum_L |
| | 3 | 29 | 3.59 | -7.65 | 12.2 | 10.9 | Dorsolateral prefrontal cortex_L |
| | 2 | 27 | 3.66 | -14.6 | 2.29 | 0.565 | Temporal area_L |
| | 1 | 27 | 4.6 | -6.66 | 20.2 | 9.92 | Dorsolateral prefrontal cortex_L |

| **$CS^-$ Activation** | Atlas Coordinates (mm) | | | | | | Brain regions |
|---|---|---|---|---|---|---|---|
| | Cluster | # Voxels | Z-Max | X | Y | Z | |
| Post-C < Post-THC | 4 | 74 | 4.37 | 4.27 | -1.69 | -7.85 | Cerebellum_R |
| | 3 | 23 | 3.82 | -3.68 | -22.6 | 2.44 | Visual cortex |
| | 2 | 22 | 3.78 | -6.66 | 19.2 | 3.37 | Orbitofrontal Cortex |
| | 1 | 21 | 4.56 | -6.66 | -13.6 | 6.18 | Parietal area_L |
| Post-C > Post-THC | N/A | | | | | | |

**Table S5.** Number of responses during Post-C and Post-THC phases. Pairwise comparisons were conducted using one-tailed paired t-tests. Statistical significance was set at *$p < 0.05$.

| | Number of responses (Mean ± SEM) | | Differences | t-ratio | p-value |
|---|---|---|---|---|---|
| | Post-C | Post-THC | | | |
| $CS^+$ sessions | 53.67 ± 5.78 | 50.67 ± 10.17 | 37.75 ± 4.36 | 0.2 | 0.43 |
